# Retinoic acid-gated BDNF synthesis in neuronal dendrites drives presynaptic homeostatic plasticity

**DOI:** 10.1101/2022.05.04.490570

**Authors:** Shruti Thapliyal, Kristin L. Arendt, Anthony G. Lau, Lu Chen

## Abstract

Homeostatic synaptic plasticity is a non-Hebbian synaptic mechanism that adjusts synaptic strength to maintain network stability while achieving optimal information processing. Among the molecular mediators shown to regulate this form of plasticity, synaptic signaling through retinoic acid (RA) and its receptor, RARα, has been shown to be critically involved in the homeostatic adjustment of synaptic transmission in both hippocampus and sensory cortices. In this study, we explore the molecular mechanism through which postsynaptic RA and RARα regulates presynaptic neurotransmitter release during prolonged synaptic inactivity at excitatory synapses. We show that RARα binds to a subset of dendritically sorted brain-derived neurotrophic factor (BDNF) mRNA splice isoforms and represses their translation. The RA-mediated translational de-repression of postsynaptic BDNF results in the retrograde activation of presynaptic Tropomyosin receptor kinase B (TrkB) receptors, facilitating presynaptic homeostatic compensation through enhanced presynaptic release. Together, our study illustrates a RA-mediated retrograde synaptic signaling pathway through which postsynaptic protein synthesis during synaptic inactivity drives compensatory changes at presynaptic site.

## Introduction

Activity in neural circuits is highly dynamic and is continuously subjected to experience-dependent changes due to synaptic plasticity. Information processing that optimally interprets the external world is vital for the survival of an organism and requires precisely timed and well-orchestrated excitatory and inhibitory synaptic transmission in neural circuits. Hebbian plasticity is considered to be the cellular mechanism underwriting memory formation. By contrast, homeostatic synaptic plasticity counters the runaway tendency of Hebbian plasticity, thus maintaining network stability while preserving its capacity for information processing. Retinoic acid (RA), a well-documented developmental morphogen, has emerged in recent studies as a critical molecular component in homeostatic synaptic plasticity [1]. Chronic synaptic inactivity at glutamatergic synapses triggers RA synthesis in neuronal dendrites, which in turn, drives local synthesis and synaptic incorporation of GluA1 containing α-amino-3-hydroxy-5-methyl-4-isoxazolepropionic acid receptors (AMPARs) and concomitant removal of γ-aminobutyric acid receptors (GABA_A_Rs) from inhibitory synapses, thus restoring synaptic excitatory/ inhibitory (E/I) balance [2-4]. The RA receptor RARα is a nuclear receptor that mediates RA-dependent transcriptional activation during development [5]. In mature neurons, however, RARα translocates out of the nucleus and acts as a molecular mediator for RA-dependent *de novo* protein synthesis [6]. In the context of homeostatic synaptic plasticity, RARα has been shown to bind to mRNAs via specific sequence motifs, and mediate the translation of these mRNAs in an RA-dependent manner. For example, activation of GluA1 synthesis upon chronic synaptic activity blockade has been demonstrated to require both RA synthesis and normal RARα expression [2, 4, 7].

Several aspects of RA-mediated homeostatic regulation of postsynaptic function have been addressed in previous studies [3, 4, 6]. Additionally, changes in presynaptic functions (i.e. release probability) as part of homeostatic synaptic mechanism have also been documented in multiple organisms [7-13]. Compared to *Drosophila* neuromuscular junctions (a type of glutamatergic synapse exhibiting robust presynaptic homeostatic plasticity), less is known about the signaling pathways involved in the homeostatic presynaptic changes at mammalian synapses. One of the classical synaptic retrograde messengers, brain-derived neurotrophic factor (BDNF), was found to be involved in the homeostatic regulation of presynaptic release in cultured hippocampal neurons [13]. Moreover, both postsynaptic AMPAR blockade-induced activation of RA signaling and the phospholipase D (PLD)-mTORC1 signaling pathway have been implicated in dendritic protein synthesis in the context of homeostatic plasticity[14, 15].

In this study, we explore whether and how synaptic RA/RARα signaling modulates postsynaptic BDNF synthesis during chronic synaptic inactivity in hippocampal pyramidal neurons. We show that RARα is directly associated with specific dendritically localized *Bdnf* mRNA isoforms. This enables RA-induced translational de-repression of BDNF synthesis in dendrites, resulting in retrograde activation of presynaptic Tropomyosin receptor kinase B (TrkB) receptors and upregulation of miniature excitatory postsynaptic current (mEPSC) frequencies. Using pre- and post-synaptic specific genetic deletions of RARα, BDNF and TrkB, we unequivocally established the locations of the actions of each of these key signaling molecules during homeostatic modulation of presynaptic function.

## Results

### Postsynaptic RA receptor RARα mediates Activity-blockade induced Presynaptic Homeostatic Plasticity

We have previously shown that in older (21 day-in-vitro, DIV) but not young (14 DIV) cultured primary hippocampal neurons, chronic treatment with postsynaptic activity blockers such as the AMPAR antagonist 6-cyano-7-nitroquinoxaline-2,3-dione (CNQX) or L-type voltage-dependent calcium channel (VDCC) inhibitor nifedipine results in a significant increase in both mEPSC frequency and mEPSC amplitude [7]. While the increase in mEPSC amplitude is observed in both young and old cultured neurons, the enhancement in frequency is only present in the older cultures [7]. Importantly, this increase in both mEPSC frequency and amplitude requires RA synthesis as pharmacological inhibition of RA synthesis blocks the increase in both parameters. Additionally, RA synthesis in postsynaptic neurons triggers presynaptic changes in a cell-autonomous manner as sparse expression of a dihydropyridine-insensitive L-VDCC, which specifically blocks RA synthesis in transfected postsynaptic neurons, prevented nifedipine-induced homeostatic increase in mEPSC frequency. However, what remains unanswered is where and through what molecular signaling pathway RA acts to trigger presynaptic changes.

To first localize RA action, we prepared organotypic hippocampal slices from RARα conditional knockout mouse [16, 17] in which Cre-mediated RARα deletion can be explicitly achieved in pre- or post-synaptic neurons, depending on the location of the expression of AAV-CRE. In uninfected WT neurons from DIV 21-25 hippocampal slice cultures, acute RA treatment (10 µM, 4 hrs) or prolonged CNQX treatment (20 µM, 36 hrs) induced a significant increase in both mEPSC amplitude and frequency in CA1 pyramidal neurons (Fig. 1A - 1C), replicating our previous findings in primary hippocampal neuronal cultures [7]. Moreover, selective deletion of RARα in postsynaptic neurons (Cre expressed in CA1 neurons), but not presynaptic neurons (Cre expressed in CA3 neurons), blocked both RA- and CNQX-induced homeostatic increases in both mEPSC amplitude and frequency (Fig. 1A – 1C). Thus, newly synthesized RA acts via postsynaptic RARα to promote the homeostatic adjustment of synaptic strength in both pre- and post-synaptic compartments.

**Figure 1.**
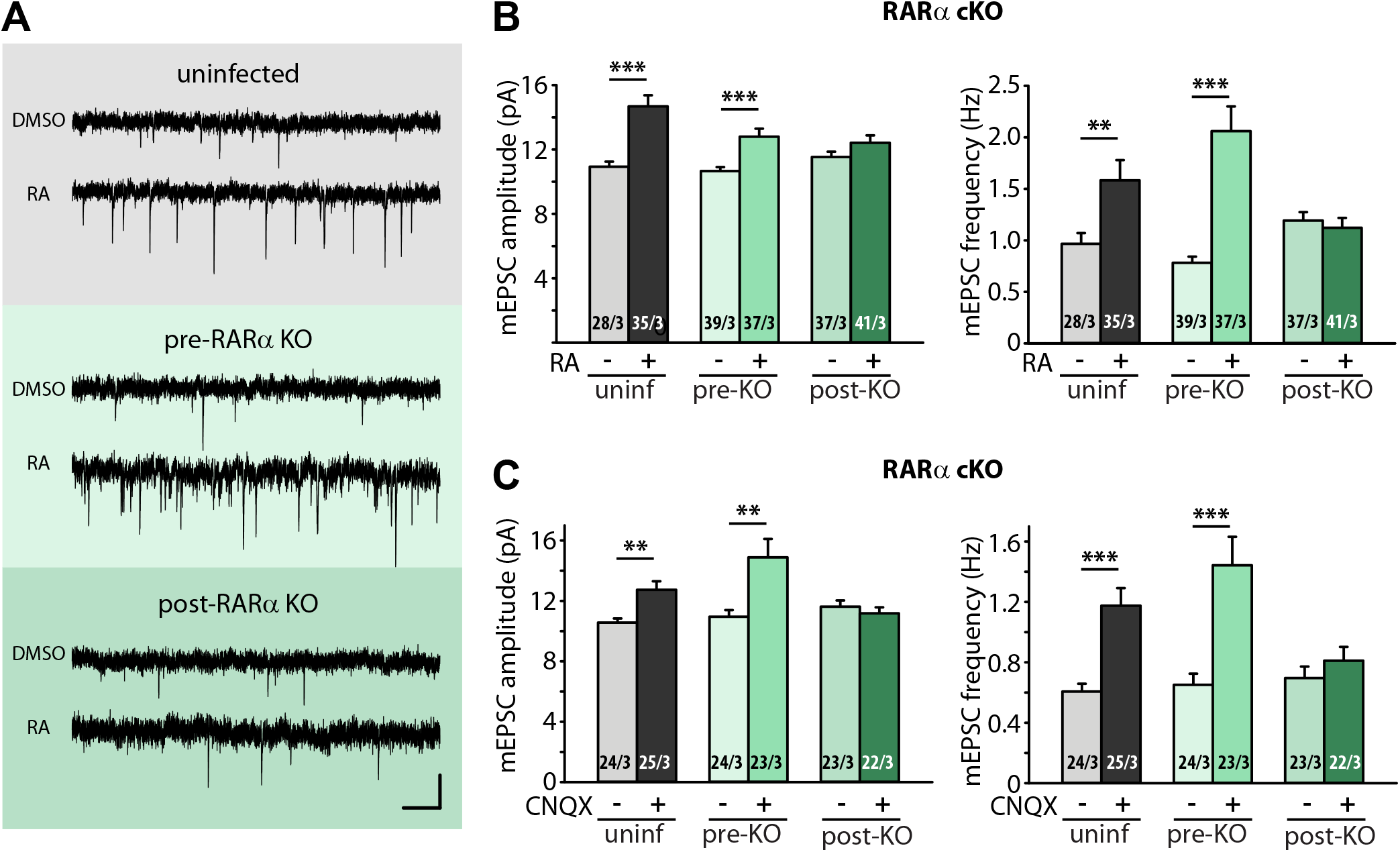
Postsynaptic RARα expression is required for presynaptic homeostatic plasticity. A) Example traces of mEPSCs recorded from hippocampal pyramidal neurons in organotypic slices from WT (uninfected), presynaptic RARα KO (Cre expression in CA3) and postsynaptic RARα KO (Cre expression in CA1) groups treated with DMSO or RA (10 µM, 4 hr). Scale bars, 10 pA, 0.5 sec. B) Quantification of mEPSC amplitudes and frequencies recorded from WT, presynaptic and postsynaptic RARα KO neurons treated with DMSO or RA. (**, *p* < 0.01; ***, *p* < 0.001; Student t-test). C) Quantification of mEPSC amplitudes and frequencies recorded from WT, presynaptic and postsynaptic RARα KO neurons treated with DMSO or CNQX (36 hours). (**, *p* < 0.01; ***, *p* < 0.001; Student t-test). n/N represent number of neurons/number of independent experiments. All graphs represent mean ± SEM.

### RA activates BDNF synthesis through direct association between RARα and specific splice isoforms of dendritically localized *Bdnf* mRNAs

A postsynaptically initiated mechanism of translational regulation mediated by RA-RARα signaling modulating presynaptic neurotransmitter release suggests the existence of a retrograde messenger molecule. Brain-derived neurotrophic factor (BDNF) is one of the most widely expressed and well-characterized neurotrophins in the developing and adult mammalian central nervous system [18, 19]. It is synthesized as a proneurotrophin known as pro-BDNF and secreted as a mixture of pro-BDNF and mature BDNF processed from pro-BDNF [20]. The retrograde BDNF-TrkB signaling has been well-established in modulating both Hebbian synaptic plasticity [21, 22] and homeostatic synaptic plasticity [13]. BDNF expression in the brain is developmentally regulated and exhibits a steep increase between postnatal day 15 and 20 [23, 24], a time period that coincides with the emergence of RA -dependent presynaptic homeostatic modulation observed in DIV21 primary cultured hippocampal neurons [7]. Immunoblotting of proBDNF in hippocampal tissue collected at different developmental stages showed a gradual increase in BDNF expression during the first four postnatal weeks (Figure 2 – figure supplement 1A). Immunoblot analysis from cultured hippocampal slices (prepared from postnatal day 8 mouse pups) showed a similar trend where the expression of BDNF gradually increases as the cultures mature from DIV1 to DIV 21 (Figure 2A), suggesting that this developmental upregulation of BDNF expression is preserved in our organotypic hippocampal slice culture system.

**Figure 2.**
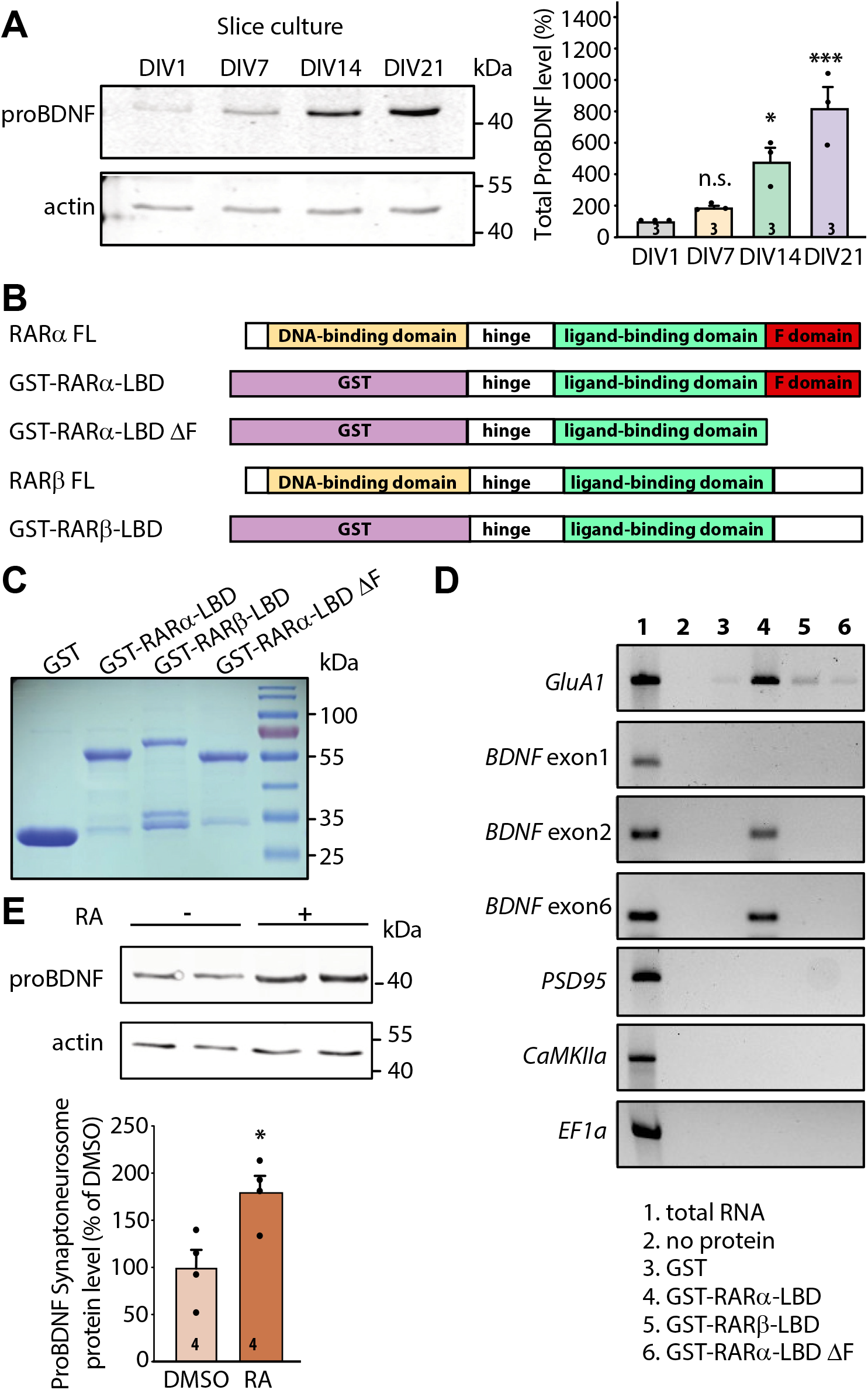
RARα binds specific *bdnf* transcript isoforms. A) Representative immunoblots (left) and quantification (right) depicting ProBDNF expression profiles in cultured hippocampal slices collected at 1, 7, 14, and 21 days in culture. Actin was used as a loading control and all expression levels were normalized to that of DIV 1 (one-way ANOVA with Dunnett’s multiple comparison test, ***, *p* < 0.001; *, p < 0.05). B) Schematic diagram of recombinant GST fusion proteins of RARα LBD, RARα LBD ΔF and RARβ LBD used in RNA binding assays. Full length RARα and RARβ protein structures are shown on as references. C) Representative imaging of Coomassie brilliant blue-stained SDS-polyacrylamide gel showing the expression of purified recombinant GST and GST fused RARα and RARβ LBD proteins (n=3). D) Representative images for semi-quantitative RT-PCR of specific BDNF transcripts pulled down from total hippocampal RNAs in *in vitro* selection with purified GST-fusion proteins. *GluA1* mRNA was used as positive control. *PSD95, CaMKIIa* and *EF1a* mRNAs served as negative controls. The representative image shown here is from one of the three experiments with similar results. E) Representative immunoblot (left) and quantification (right) showing induced ProBDNF synthesis in synaptoneurosomal fraction following 30 minutes of RA treatment. Actin was used as a loading control (two-tailed unpaired t-test, *, *p* < 0.05). N represent number of independent experiments. All graphs represent mean ± SEM.

We next sought to investigate whether postsynaptic RA/RARα is involved in retrograde BDNF signaling through regulation of BDNF synthesis. We first explored the possibility that RARα directly binds to *Bdnf* transcripts. In rodents, the *Bdnf* gene represents a complex structure that consists of nine unique promoters (on nine separate exons) that drive the expression of several distinct *bdnf* transcript isoforms (Figure 2 – figure supplement 1B). All regulatory non-coding exons yield a single identical mature BDNF protein [25]. The complex *bdnf* gene structure gives rise to a diverse array of *bdnf* transcripts that can be differentially trafficked to subcellular domains and translationally regulated by distinct intra- and extra-cellular signals [26-28]. We first performed *in silico* analysis to search for RARα recognition motifs in non-coding *Bdnf* exons. This analysis revealed that at least two *Bdnf* exons (exon 2 and exon 6) carry RARα-binding motifs in their sequence (Figure 2 – figure supplement 1B). *Bdnf* transcripts containing exon 2 or exon 6 are trafficked to distal dendrites and are subject to synaptic activity-dependent translational regulation [28-32]. In cultured hippocampal pyramidal neurons, the localization of *Bdnf* exon 2 and exon 6 in distal dendrites increases dramatically as the culture matures from DIV7 to DIV18 [29], making them potential targets for RA/RARα-mediated translational regulation around the onset time of homeostatic presynaptic changes.

RARα-mediated translational regulation in dendrites requires direct association of dendritically sorted mRNAs to RARα via specific recognition motifs [6]. Specifically, the carboxyl-terminal F domain of RARα mediates direct binding to substrate mRNAs (Figure 2B). This mRNA-binding ability is specific to RARα as another RA receptor family member RARβ, which has a different amino acid sequence in the F domain, shows only minimal association with dendritic mRNAs [6]. We first performed *in vitro* RNA binding assay in which RARα LBD domain (including the F domain) was fused to GST and immobilized on glutathione Sepharose beads. GST alone, GST-tagged RARα LBD lacking F-domain (RARα LBD ΔF), and RARβ LBD were used as negative controls. Total RNAs pooled from 3–4-week-old whole hippocampi were used in the binding assay. GST and all fusion proteins showed comparable expression and binding to the beads (Figure 2C and Figure 2 – figure supplement 1C). RT-PCR was used to test the relative enrichment of selected mRNAs by RARα. *Bdnf* transcripts containing exon 2 or exon 6 exhibited selective enrichment through binding to RARα LBD, along with GluA1 mRNAs, a mRNA substrate of RARα identified in our previous study [6] (Figure 2D). Transcripts of PSD95, CaMKII and EF1α, which are also dendritically localized but do not exhibit RARα binding in previous studies [6], served as negative controls (Figure 2D). By contrast, *Bdnf* transcript containing exon 1 that carries only a single RARα recognition motif (Figure 2 – figure supplement 1B) and is not sorted to dendrites in hippocampal neurons [29], failed to bind to RARα LBD (Figure 2D). Additionally, deletion of F domain, the mRNA-binding domain at the C-terminal end of RARα, abolished the binding activity of all mRNAs (Figure 2D). Taken together, this data indicates that the two dendritically localized *Bdnf* mRNAs (carrying exon 2 and exon 6) exhibit specific binding to RARα F domain.

Does the binding of *Bdnf* mRNAs by RARα result in their translational regulation by RA? To address this, we examined RA-induced local translation of BDNF in synaptoneurosomes (SNS) prepared from 3-4 weeks old mouse hippocampi. The purity of synaptoneurosomal fraction was verified by the enrichment of synaptic protein PSD95 and the absence of nuclear protein Histone H3 (Figure 2 – figure supplement 1D). A brief treatment of synaptoneurosomes with 1 μM RA for 30 minutes at 37°C significantly increased GluA1 protein levels and demonstrated the translational competency of the synaptoneurosomal preparation [2] (Figure 2 – figure supplement 1E). Importantly, the expression levels of ProBDNF protein also increased significantly (Figure 2E) while PSD95, whose mRNAs are also dendritically localized but do not bind to RARα, failed to respond to RA stimulation [6](Figure 2 – figure supplement 1F). Thus, RARα binding to *Bdnf* mRNAs does convey translational regulation of BDNF by RA.

### RA drives retrograde BDNF-TrkB signaling to achieve regulation of presynaptic homeostatic plasticity

Having established that RA regulates BDNF synthesis through direct binding between RARα and dendritically localized isoforms of *Bdnf* mRNAs, we next sought to test the relevance between RA-dependent BDNF synthesis and presynaptic homeostatic changes. To this end, we prepared organotypic hippocampal slice cultures from conditional knockout mice for either BDNF or TrkB, which allowed for dissection of the loci of action (pre vs postsynaptic) for these two key players. Injection of Cre-expressing AAVs into either the CA3 or CA1 regions of the hippocampus achieved pre- or postsynaptic-specific deletion of target proteins in the Schaeffer collateral-CA1 synapses. Acute RA treatment or prolonged CNQX treatment significantly increased both mEPSC amplitudes and frequencies in uninfected control slices (Figures 3A-3C). While presynaptic deletion of BDNF did not impair either of these changes in mEPSCs, postsynaptic deletion of BDNF prevented the increase in mEPSC frequency (Figures 3B-3C). By contrast, presynaptic, but not postsynaptic deletion of TrkB blocked the increase in frequency induced by either RA or CNQX (Figures 3D-3F). Deletion of either BDNF or TrkB pre or postsynaptically did not affect the homeostatic increase in mEPSC amplitude, indicating that BDNF/TrkB signaling is exclusively involved in regulation of presynaptic function. The locations of action for BDNF and TrkB are consistent with the postsynaptic initiation of RA synthesis followed by RARα-mediated translational regulation supporting the notion that retrograde BDNF signaling through presynaptic TrkB drives presynaptic changes during homeostatic synaptic plasticity (Figure 3 – figure supplement 1).

**Figure 3.**
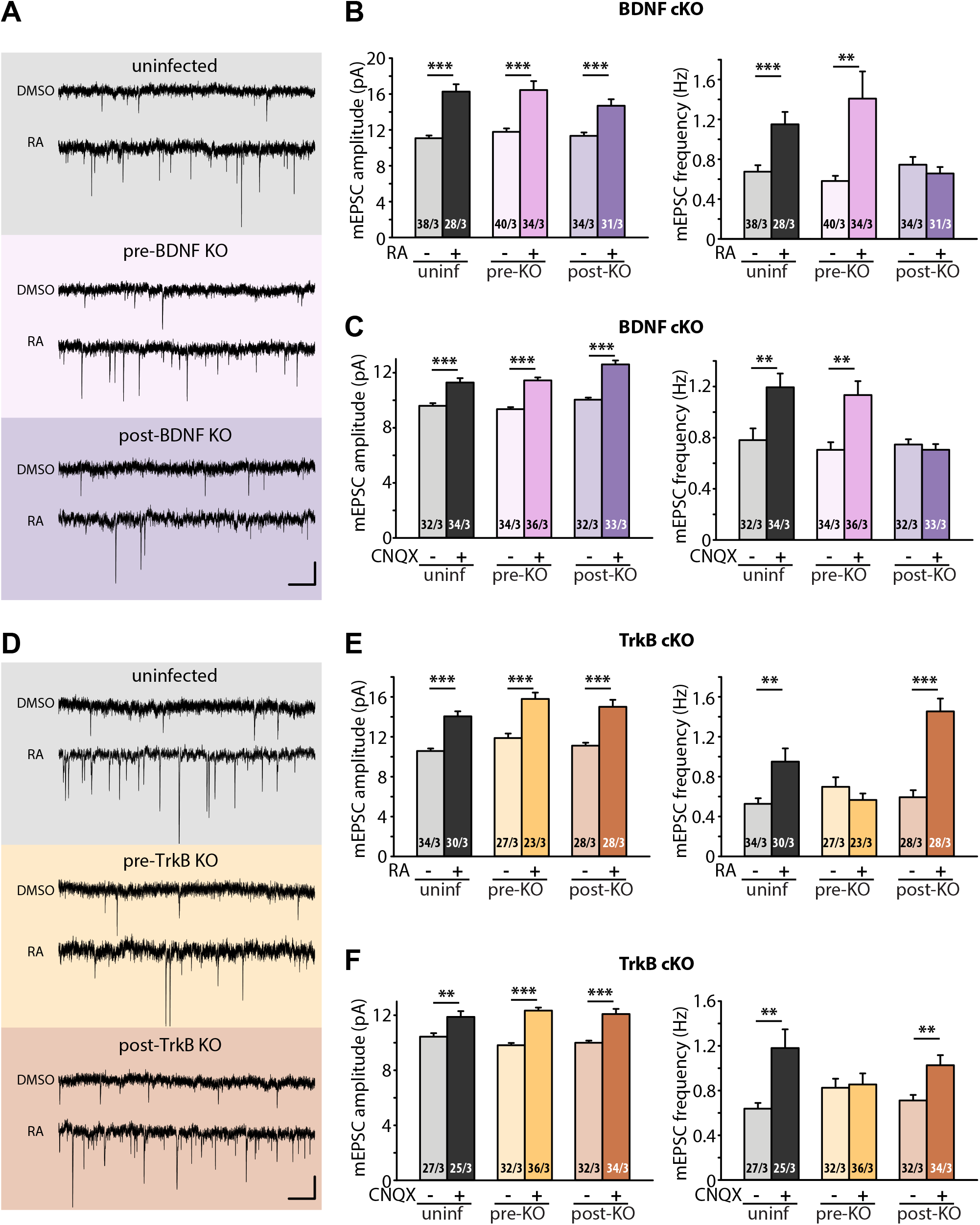
Retrograde BDNF signaling is required for RARα-mediated regulation of presynaptic homeostatic scaling. A) Example traces of mEPSCs recorded from hippocampal pyramidal neurons in organotypic slices from WT (uninfected), presynaptic BDNF KO (Cre expression in CA3) and postsynaptic BDNF KO (Cre expression in CA1) groups treated with DMSO or RA (10 µM, 4 hr). Scale bars, 10 pA, 0.5 sec. B) Quantification of mEPSC amplitudes and frequencies recorded from WT, presynaptic and postsynaptic BDNF KO neurons treated with DMSO or RA. (**, *p* < 0.01; ***, *p* < 0.001; Student t-test). C) Quantification of mEPSC amplitudes and frequencies recorded from WT, presynaptic and postsynaptic BDNF KO neurons treated with DMSO or CNQX (36 hours). (**, *p* < 0.01; ***, *p* < 0.001; Student t-test). D) Example traces of mEPSCs recorded from hippocampal pyramidal neurons in organotypic slices from WT (uninfected), presynaptic TrkB KO (Cre expression in CA3) and postsynaptic TrkB KO (Cre expression in CA1) groups treated with DMSO or RA (10 µM, 4 hr). Scale bars, 10 pA, 0.5 sec. E) Quantification of mEPSC amplitudes and frequencies recorded from WT, presynaptic and postsynaptic TrkB KO neurons treated with DMSO or RA. (**, *p* < 0.01; ***, *p* < 0.001; Student t-test). F) Quantification of mEPSC amplitudes and frequencies recorded from WT, presynaptic and postsynaptic TrkB KO neurons treated with DMSO or CNQX (36 hours). (**, *p* < 0.01; ***, *p* < 0.001; Student t-test). n/N represent number of neurons/number of independent experiments. All graphs represent mean ± SEM.

## Discussion

This study describes an RA/RARα-dependent trans-synaptic retrograde signaling pathway that modulates presynaptic function during homeostatic plasticity. Here, we identify dendritically localized postsynaptic *Bdnf* transcripts as RARα-binding targets that are subject to RA-dependent regulation of BDNF synthesis upon activity blockade. BDNF enhances presynaptic function via a retrograde signaling mechanism by binding to presynaptic TrkB receptors. In concert with an RA-dependent increase in postsynaptic AMPA receptor abundance, this molecular pathway contributes to homeostatic adjustment of synaptic strength during chronic activity blockade.

BDNF has diverse roles in synapse function and plasticity and is implicated in the pathophysiology of various brain disorders [33-37]. As one of the key trans-synaptic signaling molecules, the transcription, synthesis, and secretion of BDNF are highly activity-dependent [38-42]. This activity-dependent property of BDNF expression and secretion plays important roles in synapse development and remodeling [43-51]. Moreover, through its actions on both inhibitory and excitatory synapses, BDNF signaling impacts synaptic function [52-56], activity-dependent synaptic plasticity mechanisms such as long-term potentiation (LTP) and long-term depression (LTD) [57-63], and ultimately learning and memory formation [64-68]. However, how synaptic inactivity regulates BDNF expression in the context of homeostatic synaptic plasticity is not yet fully understood.

The impact of exogenous BDNF on basal synaptic function and its connection to homeostatic plasticity has been investigated in several studies [26, 69]. In visual cortical pyramidal neurons, exogenous BDNF blocks homeostatic upscaling induced by chronic synaptic inactivity while BDNF depletion with TrkB-IgG scales up mEPSC amplitudes [70]. While these results suggest a potential involvement of BDNF signaling in cortical pyramidal neuron homeostatic plasticity, chronic treatment of BDNF does not downscale mEPSCs [71]. Moreover, BDNF’s effect on basal synaptic transmission seems to be cell type- and region-specific. Different from its effect on cortical pyramidal cells, BDNF increases mEPSC amplitudes in visual cortical interneurons [70]. In the hippocampus, BDNF enhances mEPSC amplitudes without affecting their frequencies, but increases mIPSC frequency and sizes of GABAergic synaptic terminals [72]. In nucleus accumbens medium spiny neurons (MSNs), acute BDNF treatment (30 min) increases surface AMPA receptor expression, but chronic BDNF treatment (24 hr) decreases surface AMPA receptor expression and blocks synaptic downscaling [73, 74]. Taken together, it becomes apparent that BDNF signaling is complex as its effect on synaptic transmission is multifaceted and dependent on the context of the study (cell type, brain region, developmental stage, etc.,) [26].

In addition to studies examining the impact of exogenous BDNF on basal synaptic transmission, the involvement of endogenous BDNF signaling in homeostatic synaptic plasticity has been further investigated and found to be required for both synaptic upscaling and downscaling in hippocampal neurons [8, 13, 42]. In the case of synaptic hyperactivity-induced synaptic downscaling, mIPSC blockade and postsynaptic increase in calcium influx were found to induce postsynaptic increase in *Bdnf* transcription, which mediates homeostatic down-regulation of mEPSC amplitudes [42]. Paradoxically, in the case of synaptic upscaling, postsynaptic release of BDNF as a retrograde messenger was also required in synaptic activity blockade-induced homeostatic increase in presynaptic vesicle turnover [8] and mEPSC frequency [13]. Furthermore, rapid activation of dendritic mTORC1 signaling was proposed to be a molecular mechanism for protein-synthesis dependent release of postsynaptic BDNF to mediate presynaptic homeostatic compensation under prolonged inactivation of postsynaptic AMPA receptors[14, 15].

How does the same synaptic signaling molecule (BDNF) act in seemingly opposite biological processes? The vastly diverse functions of BDNF in the nervous system have been attributed to tight spatial, temporal, and stimulus-dependent regulation of BDNF expression from its complex gene structure. Using a combination of individual promoters (served by initial 8 non-coding exons in rodents) and polyadenylation sites, a single *Bdnf* gene could give rise to several different transcripts [25]. Although it is widely accepted that most of these individual *Bdnf* promoters respond differentially to neuronal activity, the specific functional roles of most of these transcripts remain elusive. For instance, multiple studies have shown that promoter/exon IV is robustly involved in activity-dependent BDNF expression in the cortex [75-80]. In addition, BDNF expression driven by promoter/exon I, III and VI has been shown to be regulated by AP1 transcription factors and by TrkB signaling in a positive feedback loop [53, 81]. However, how synaptic inactivity modulates specific-exon derived BDNF expression and which regulatory exons/promoters of the *Bdnf* gene respond to synaptic inactivity has never been explored. Further, how specific *Bdnf* transcripts contribute to homeostatic scaling is largely unknown. Our study focuses on the involvement of different *Bdnf* transcripts in synaptic inactivity-induced presynaptic upscaling. We found that dendritically localized *Bdnf* transcripts containing exon II and VI are subjected to RA/RARα-mediated regulation of synthesis in the postsynaptic compartment under chronic synaptic activity blockade. These newly synthesized BDNF acts retrogradely on presynaptic TrkB receptors to mediate homeostatic modulation of presynaptic function. An increase in *Bdnf* transcription and enhanced dendritic targeting of specific *Bdnf* transcripts have been associated with increased neuronal and synaptic activity [40, 42, 82-86]. Our study shows that by acting on specific dendritically targeted *Bdnf* transcripts, RA/RARα selectively activates BDNF synthesis in postsynaptic compartments during synaptic inactivity, thus further extending the molecular regulatory mechanisms underlying BDNF signaling in synaptic inactivity-induced synaptic plasticity.

Similar to the rodent *Bdnf* gene, the human *BDNF* gene has a complex structure with multiple transcription initiation sites and alternative transcripts [87], and is implicated in neuropsychiatric disorders [26, 37, 88]. The involvement of synaptic RA/RARα signaling in various neuropsychiatric diseases has begun to emerge in recent years [89-92]. In particular, the absence of RA-RARα regulated homeostatic plasticity mechanisms have been established in both *Fmr1* knockout mice (a mouse model for fragile X syndrome (FXS)) and human neurons differentiated from FXS patient-derived induced pluripotent stem cells [93-95]. The convergence of BDNF signaling and RA signaling for synaptic functions and homeostatic plasticity highlights the importance of understanding the intersections of various synaptic signaling pathways and their implications in cognitive function.

## Author Contributions

All authors participated in experiment design and analysis. S.T., K.L.A. and A.G.L. performed all the experiments and analyses with input from L.C.. S.T. and L.C. wrote the manuscript with input from K.L.A..

## Acknowledgments

We thank Drs. Xiling Li, Michelle Tjia and Omid Miry for their helpful input and technical support. The work was supported by NIH grants MH086403 (L.C.), NS11566001 (L.C.), HD104458 (L.C.).

## Supplemental Information

Supplemental Information includes three figures, which can be found with this article online at <URL>.

## Material and Methods

### Key Reagents and Resources table

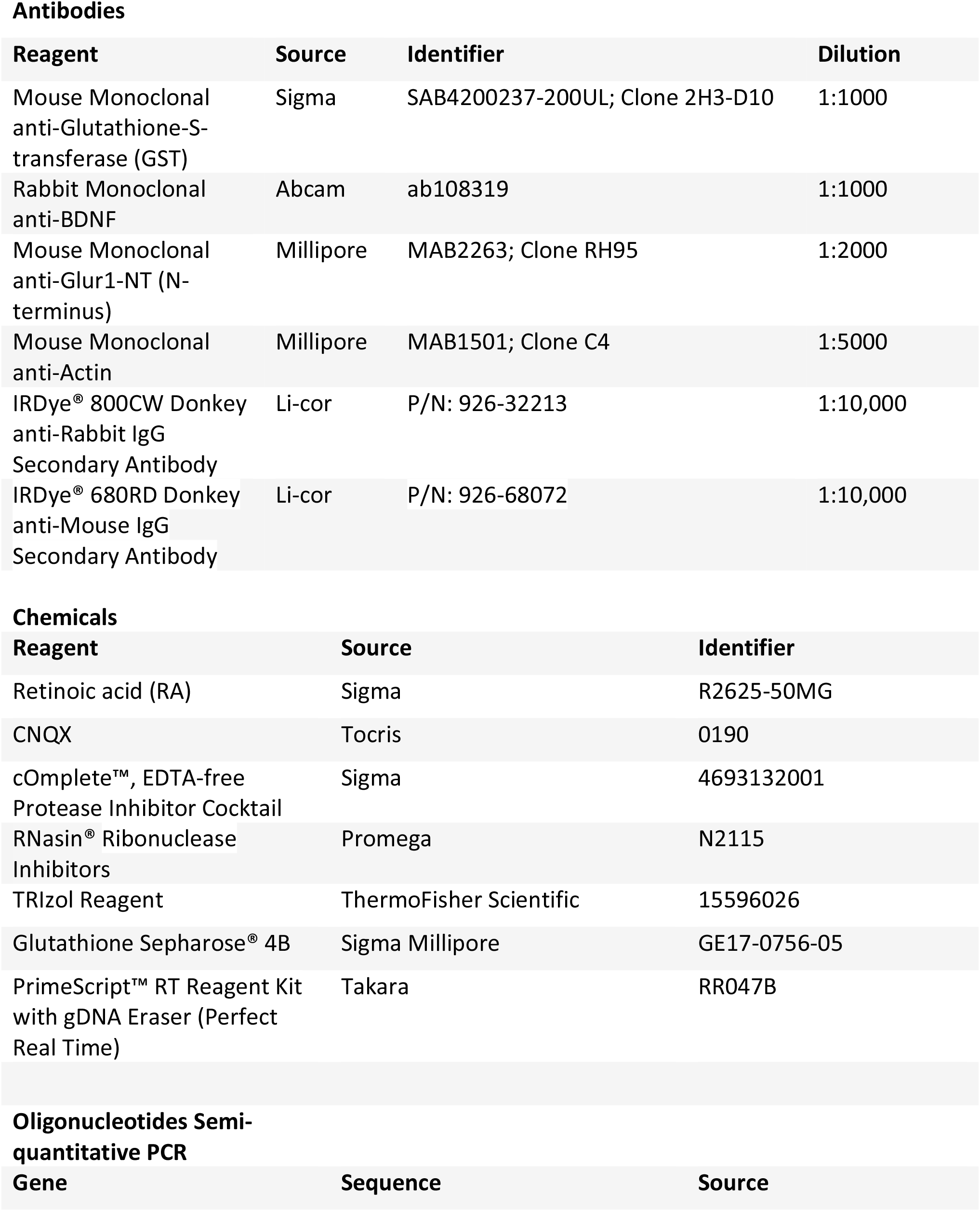
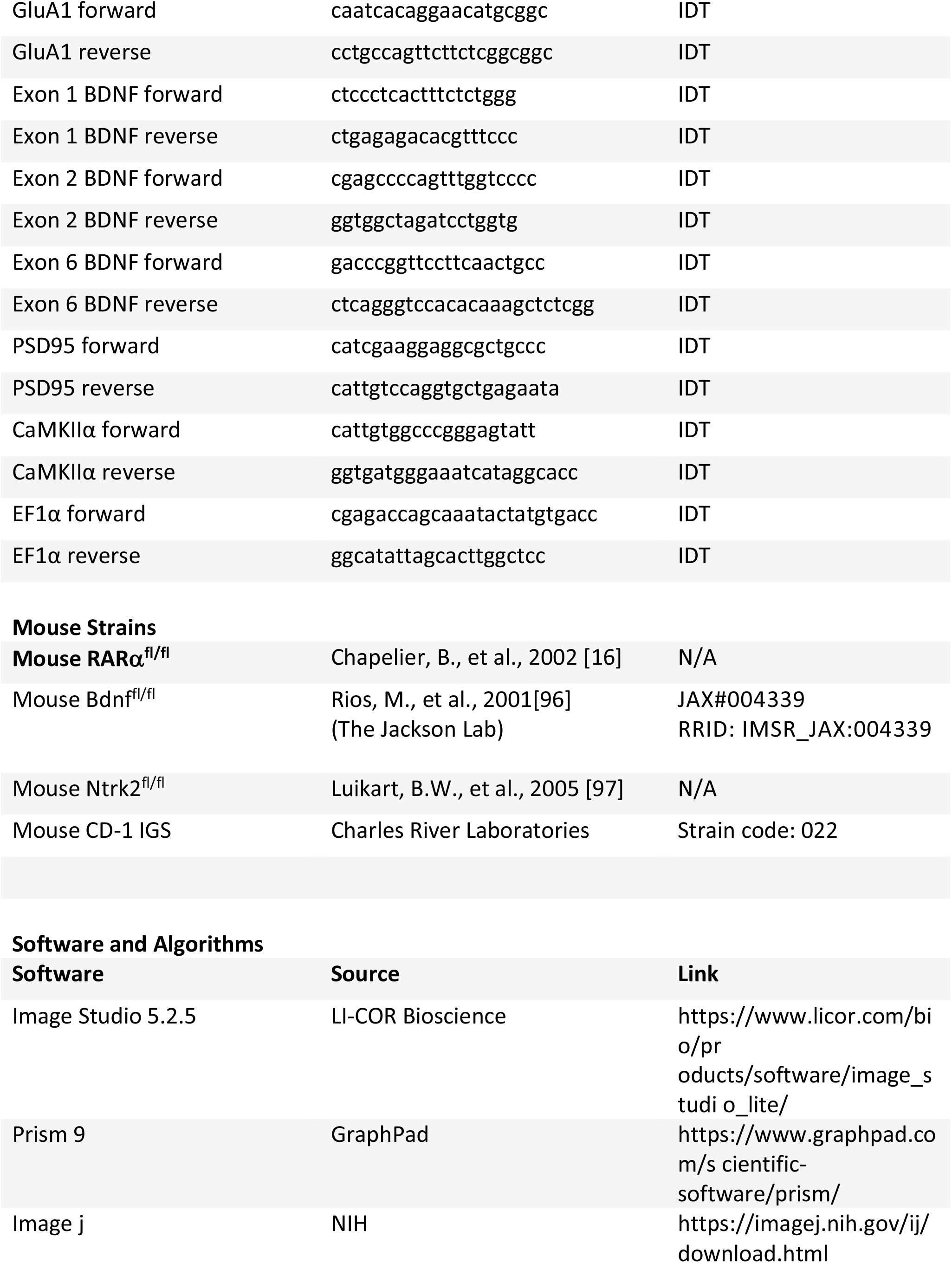
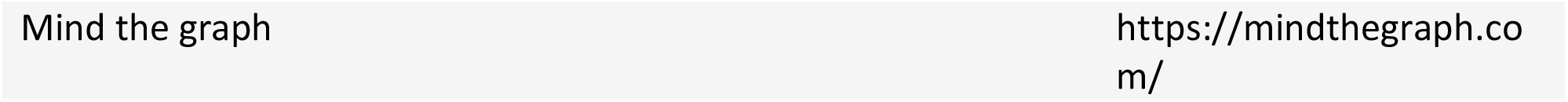

### Animals

Mouse strains used in the study are mentioned in table above. All mouse studies were performed according to protocols approved by the Stanford University Administrative Panel on Laboratory Animal Care. All procedures conformed to NIH Guidelines for the Care and Use of Laboratory Animals and were approved by the Stanford University Administrative Panel.

### Plasmid constructs for Recombinant gene expression

All RAR constructs were generated using mouse sequences and cloned into pGEX-KG. For GST-RARα LBD, nucleotides 460 to end were cloned using BamHI and HindIII; for GST-RARα LBD ΔF, nucleotides 460 to 1251 were cloned with BamHI and HindIII; and for GST-RARβ LBD, nucleotides 496 to end were cloned using BamHI and HindIII.

### *In-vitro* RNA binding assay

Purified GST fusion proteins and total RNA from whole hippocampi of P25-P30 mouse pups were used for selection. GST fusion proteins were expressed in BL21 cells induced with 1 mM IPTG. Bacteria was sonicated in lysis buffer (150 mM NaCl, 20 mM sodium phosphate, pH 7.4, 1% Triton X-100, and protease inhibitors) and debris were cleared by centrifugation at 10,000 rpm for 20 minutes. Protein expression was confirmed by SDS/PAGE followed by Coomassie staining and immunoblotting when possible. GST fused to the various domains of RARα or RARβ were then purified from the bacterial lysate by binding to glutathione Sepharose beads and equilibrating/washing the protein-bound beads five times in RNA binding buffer (200 mM KOAc, 10 mM TrisOAc, pH 7.7, and 5 mM MgOAc with protease and RNase inhibitors). Total RNA was obtained from whole hippocampi of 25–30-day-old CD1 mice using TRIzol. RNA was DNase treated and reextracted with TRIzol, and the pellet was re-suspended in nuclease-free water and quantified by spectrophotometry. RNA (20 μg) was added to RNA binding buffer and heated to 95 °C to denature secondary structure, then slowly re-natured. Renatured RNA was then added to the immobilized GST-fusion protein in RNA binding buffer and rotated overnight at 4 °C. Beads were then washed several times in RNA binding buffer. RNA was extracted with TRIzol, treated with RNase free DNase I, then reverse transcribed with oligo(dT) according to manufacturer’s instructions. cDNA was used for amplification with PCR using gene-specific primers.

### Synaptoneurosome preparation

Hippocampi from P25-P30 CD1 mice were dissected and gently homogenized in a solution containing 33% sucrose, 10 mM HEPES, 0.5 mM EGTA (pH 7.4), and protease inhibitors. Nuclei and other debris were pelleted at 2,000 g for 5 min at 4°C and the supernatant was filtered through three layers of 100 μm pore nylon mesh (Millipore), and a 5 μm pore PVDF syringe filter (Millipore). The filtrate was then centrifuged for 10 min at 10,000 g at 4°C and the supernatant was removed. The synaptoneurosome-containing pellet was then resuspended in the appropriate amount of lysis buffer containing 140 mM NaCl, 3 mM KCl, 10 mM Glucose, 2 mM MgSO_4,_ 2 mM CaCl_2_ and 10 mM HEPES (pH 7.4) containing protease inhibitors and RNAsin. Equal volumes were then aliquoted into opaque microfuge tubes. Appropriate samples were incubated with 1 μM RA for 10 min at 37°C and immediately frozen in dry ice afterwards.

### Western blotting

Samples were run on 10% SDS/PAGE and transferred to nylon membranes. Membranes were blocked with Tris-buffered saline solution containing 0.1% Tween-20 [TBST] and 5% dry milk. Primary antibodies were diluted into TBST and incubated overnight at 4 °C. Primary antibody was washed with TBST and secondary antibody was added for 1 h in TBST. Secondary antibody was washed off with TBST and signal detected using Odyssey CLx imaging system (LI-COR). The densitometric analysis was performed using ImageJ software.

### Organotypic hippocampal slice cultures

Organotypic slice cultures were prepared from postnatal day 7 to 8 mouse pups and placed on semiporous membranes (Milipore) for 21-25 days prior to recording (B. H. Gahwiler et al., 37 1997). Briefly, slices were maintained in a MEM based culture media comprised of 1 mM CaCl2, 2 mM MgSO4, 1 mM L-glutamine, 1mg/L insulin, 0.0012% ascorbic acid, 30mM HEPES, 13 mM D-glucose, and 5.2 mM NaHCO3. Culture media was a pH of 7.25 and the osmolarity was 320. Cultures were maintained in an incubator with 95% O2/ 5% CO2 at 34 degrees C.

### Viral vectors and viral infection

AAV Syn-CRE was prepared as previously described [98]. Viral titers were determined by qPCR.

Cultures were injected on DIV0 and maintained for 21-25 days prior to recording. Presynaptic deletion was accomplished via injection of AAV-CRE into the CA3 pyramidal cell body layer. Postsynaptic deletion was achieved via injection of AAV-CRE into the CA1 pyramidal cell body layer. All experiments are executed with interweaving controls (either uninfected (i.e. WT), presynaptic deletion, or postsynaptic deletion) All injections were verified and confirmed as 95-100% infectivity in either the CA3 or CA1 prior to recording.

### Electrophysiology

Voltage-clamp whole-cell recordings are obtained from CA1 pyramidal neurons treated with either vehicle controls,10 uM RA for 4 hours prior to recording, or 20 uM CNQX for 36 hours prior to recording, under visual guidance using transmitted light illumination. Vehicle control, RA treated, and CNQX treated cells were obtained from the same batches of slices on the same experimental day.

The recording chamber is perfused with 119 mM NaCl, 2.5 mM KCl, 4 mM CaCl2, 4 mM MgCl2, 26 mM NaHCO3, 1 mM NaH2PO4, 11 mM glucose, and 0.1 mM picrotoxin, at pH 7.4, gassed with 5% CO2/95% O2 and held at 30ºC. Patch recording pipettes (3–6 M#) are filled with 115 mM cesium methanesulfonate, 20 mM CsCl, 10 mM HEPES, 2.5 mM MgCl2, 4 mM Na2ATP, 0.4 mM Na3GTP, 10 mM sodium phosphocreatine, and 0.6 mM EGTA at pH 7.25.

Spontaneous miniature transmission was obtained in the presence of 1uM of TTX in the external solution. For slices previously exposed to RA or CNQX, slices were washed out prior to recording spontaneous responses.

### Statistical analyses

All graphs represent average values ± s.e.m. Statistical differences were calculated according to parametric or nonparametric tests (indicated in figure legends). Comparisons between multiple groups were performed either with one-way ANOVA or with the Kruskal-Wallis ANOVA. When significant differences were observed, p values for pairwise comparisons were calculated according to two-tailed Mann-Whitney tests (for unpaired data) or Wilcoxon tests (for paired data). Comparisons between cumulative distributions were performed according to two-sample Kolmogorov–Smirnov tests. p values are indicated in each figure.

---

**Figure 2– figure supplement 1.**
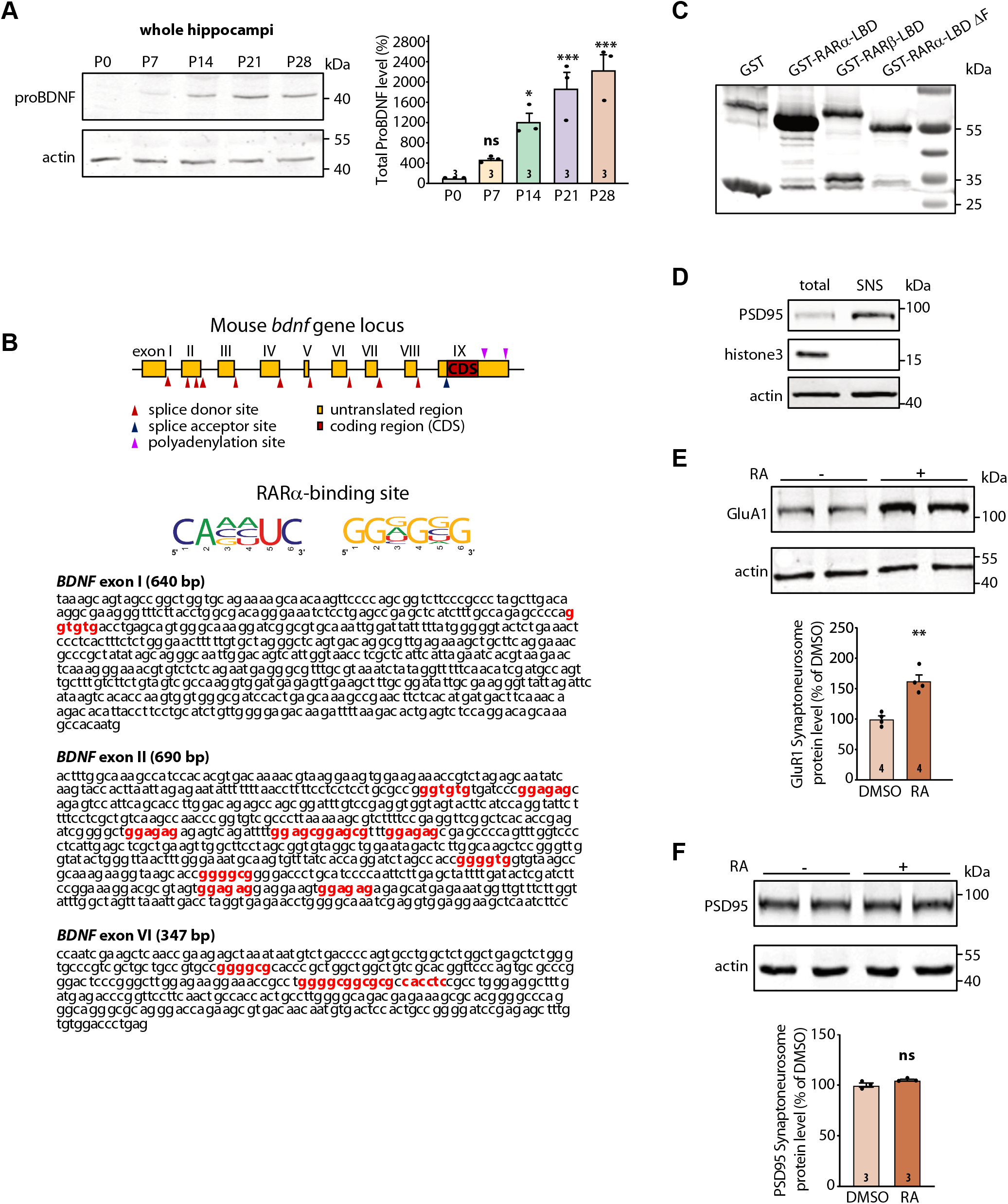
Additional data related to Figure 2: Mouse *bdnf* gene structure and expression; local translation of specific proteins induced by RA. A) Representative immunoblots (left) and quantification (right) depicting ProBDNF expression profile in whole hippocampi collected from 1-, 7-, 14-, 21- and 28-day old mouse pups. Actin was used as a loading control. All expression levels were normalized to that of P0 (one-way ANOVA with Dunnett’s multiple comparison test, ***, *p* < 0.001; *, *p* < 0.05). Graph represents mean ± SEM. B) Schematic diagram of mouse *bdnf* gene structure (top). Exons are indicated by boxes. Maroon box inside exon IX represents the coding region of the *bdnf* gene. Red arrow heads indicate the alternative splice donor sites giving rise to *bdnf* transcripts with different exon choices. The middle panel shows the specific mRNA sequence consensus recognized by RARα. The bottom panel indicates the presence of RARα-binding motifs (in red) in the 5’UTRs of three BDNF transcripts with exons I, II and VI. C) Representative GST-immunoblot showing the expression levels of purified recombinant GST and GST-tagged RARα and RARβ LBD proteins (n = 2). D) Representative immunoblots to confirm the purity of synaptoneurosome preparation. PSD95 was enriched in, and histone H3 was selectively absent from, the hippocampal synaptoneurosome fraction relative to the whole-cell lysate. E) Representative immunoblot (top) and quantification (bottom) showing induced GluA1 synthesis in synaptoneurosomal fraction following 30 minutes of RA (1 µM) treatment. Actin was used as a loading control (two-tailed unpaired t-test, **, *p* < 0.005). F) Representative immunoblot (top) and quantification (bottom) showing no induction of PSD95 synthesis in synaptoneurosomes following 30 minutes of RA treatment. Actin was used as a loading control. All graphs represent mean ± SEM.

**Figure 3– figure Supplement 1.**
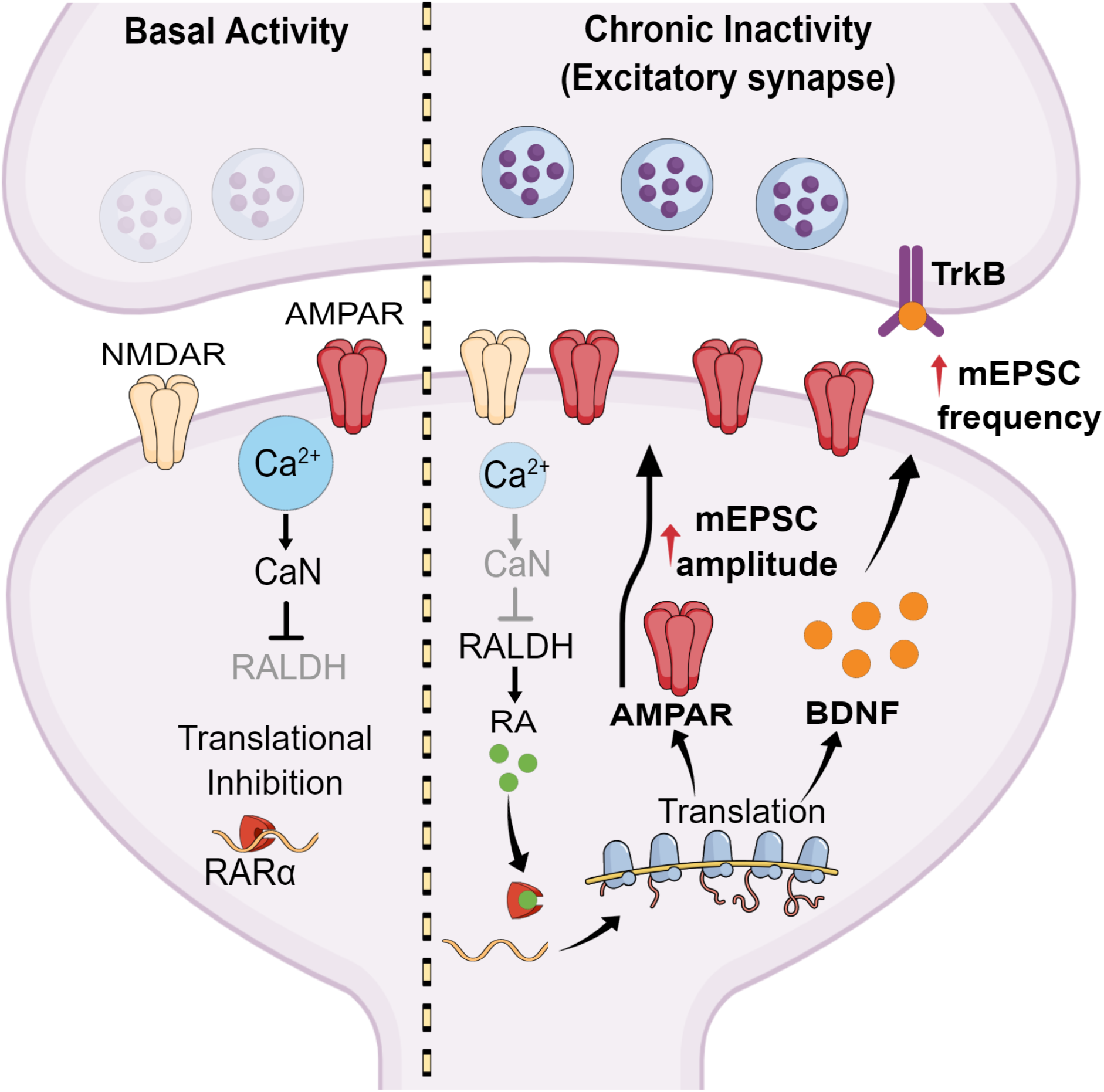
A working model for RA-mediated regulation of presynaptic and postsynaptic homeostatic plasticity. During normal synaptic transmission, basal dendritic calcium levels inhibit RA synthesis through a calcineurin-dependent pathway. This allows RARα to inhibit translation of specific dendritic mRNAs. During chronic synaptic inactivity, decreased dendritic calcium levels removes the inhibition on RA synthesis. Newly synthesized RA binds to RARα and de-represses local translation of specific dendritic mRNAs, which results in homeostatic pre- and post-synaptic changes. Among these, newly synthesized homomeric GluA1 AMPA receptors are inserted into local synapses to enhance excitatory postsynaptic strength. Additionally, newly synthesized BDNF is released from the postsynaptic neuron, acts on its presynaptic receptor TrkB, and increases presynaptic function.

## References

1. Chen, L., A.G. Lau, and F. Sarti, Synaptic retinoic acid signaling and homeostatic synaptic plasticity. Neuropharmacology, 2014. 78: p. 3–12.

2. Aoto, J., et al., Synaptic signaling by all-trans retinoic acid in homeostatic synaptic plasticity. Neuron, 2008. 60(2): p. 308–20.

3. Sarti, F., et al., Rapid suppression of inhibitory synaptic transmission by retinoic acid. J Neurosci, 2013. 33(28): p. 11440–50.

4. Maghsoodi, B., et al., Retinoic acid regulates RARalpha-mediated control of translation in dendritic RNA granules during homeostatic synaptic plasticity. Proc Natl Acad Sci U S A, 2008. 105(41): p. 16015–20.

5. Chambon, P., A decade of molecular biology of retinoic acid receptors. FASEB journal : official publication of the Federation of American Societies for Experimental Biology, 1996. 10(9): p. 940–54.

6. Poon, M.M. and L. Chen, Retinoic acid-gated sequence-specific translational control by RARalpha. Proc Natl Acad Sci U S A, 2008. 105(51): p. 20303–8.

7. Wang, H.L., et al., Decrease in calcium concentration triggers neuronal retinoic acid synthesis during homeostatic synaptic plasticity. J Neurosci, 2011. 31(49): p. 17764–71.

8. Lindskog, M., et al., Postsynaptic GluA1 enables acute retrograde enhancement of presynaptic function to coordinate adaptation to synaptic inactivity. Proceedings of the National Academy of Sciences of the United States of America, 2010. 107(50): p. 21806–11.

9. Thiagarajan, T.C., M. Lindskog, and R.W. Tsien, Adaptation to synaptic inactivity in hippocampal neurons. Neuron, 2005. 47(5): p. 725–737.

10. Davis, G.W. and M. Muller, Homeostatic control of presynaptic neurotransmitter release. Annual review of physiology, 2015. 77: p. 251–70.

11. Murthy, V.N., et al., Inactivity produces increases in neurotransmitter release and synapse size. Neuron, 2001. 32(4): p. 673–82.

12. Gong, B., et al., Genetic evidence for the requirement of adenylyl cyclase 1 in synaptic scaling of forebrain cortical neurons. The European journal of neuroscience, 2007. 26(2): p. 275–88.

13. Jakawich, S.K., et al., Local presynaptic activity gates homeostatic changes in presynaptic function driven by dendritic BDNF synthesis. Neuron, 2010. 68(6): p. 1143–58.

14. Henry, F.E., et al., Retrograde changes in presynaptic function driven by dendritic mTORC1. J Neurosci, 2012. 32(48): p. 17128–42.

15. Henry, F.E., et al., A Unique Homeostatic Signaling Pathway Links Synaptic Inactivity to Postsynaptic mTORC1. J Neurosci, 2018. 38(9): p. 2207–2225.

16. Chapellier, B., et al., A conditional floxed (loxP-flanked) allele for the retinoic acid receptor alpha (RARalpha) gene. Genesis (New York, N.Y.: 2000), 2002. 32(2): p. 87–90.

17. Sarti, F., et al., Conditional RARalpha knockout mice reveal acute requirement for retinoic acid and RARalpha in homeostatic plasticity. Front Mol Neurosci, 2012. 5: p. 16.

18. Hofer, M., et al., Regional distribution of brain-derived neurotrophic factor mRNA in the adult mouse brain. The EMBO journal, 1990. 9(8): p. 2459–64.

19. Conner, J.M., et al., Distribution of brain-derived neurotrophic factor (BDNF) protein and mRNA in the normal adult rat CNS: Evidence for anterograde axonal transport. Journal of Neuroscience, 1997. 17(7): p. 2295–2313.

20. Foltran, R.B. and S.L. Diaz, BDNF isoforms: a round trip ticket between neurogenesis and serotonin? Journal of neurochemistry, 2016. 138(2): p. 204–21.

21. Andero, R., D.C. Choi, and K.J. Ressler, BDNF-TrkB receptor regulation of distributed adult neural plasticity, memory formation, and psychiatric disorders. Progress in molecular biology and translational science, 2014. 122: p. 169–92.

22. Guo, W., G. Nagappan, and B. Lu, Differential effects of transient and sustained activation of BDNF-TrkB signaling. Developmental neurobiology, 2018. 78(7): p. 647–659.

23. Schoups, A.A., et al., NGF and BDNF are differentially modulated by visual experience in the developing geniculocortical pathway. Brain research. Developmental brain research, 1995. 86(1-2): p. 326–34.

24. Dincheva, I., N.B. Lynch, and F.S. Lee, The Role of BDNF in the Development of Fear Learning. Depression and anxiety, 2016. 33(10): p. 907–916.

25. Aid, T., et al., Mouse and rat BDNF gene structure and expression revisited. Journal of neuroscience research, 2007. 85(3): p. 525–35.

26. Wang, C.S., E.T. Kavalali, and L.M. Monteggia, BDNF signaling in context: From synaptic regulation to psychiatric disorders. Cell, 2022. 185(1): p. 62–76.

27. Lau, A.G., et al., Distinct 3’UTRs differentially regulate activity-dependent translation of brain-derived neurotrophic factor (BDNF). Proc Natl Acad Sci U S A, 2010. 107(36): p. 15945–50.

28. Song, M., K. Martinowich, and F.S. Lee, BDNF at the synapse: why location matters. Molecular psychiatry, 2017. 22(10): p. 1370–1375.

29. Baj, G., et al., Spatial segregation of BDNF transcripts enables BDNF to differentially shape distinct dendritic compartments. Proceedings of the National Academy of Sciences of the United States of America, 2011. 108(40): p. 16813–8.

30. Colliva, A. and E. Tongiorgi, Distinct role of 5’UTR sequences in dendritic trafficking of BDNF mRNA: additional mechanisms for the BDNF splice variants spatial code. Molecular brain, 2021. 14(1): p. 10.

31. Vaghi, V., et al., Pharmacological profile of brain-derived neurotrophic factor (BDNF) splice variant translation using a novel drug screening assay: a “quantitative code”. The Journal of biological chemistry, 2014. 289(40): p. 27702–13.

32. Baj, G., et al., Signaling pathways controlling activity-dependent local translation of BDNF and their localization in dendritic arbors. Journal of cell science, 2016. 129(14): p. 2852–64.

33. Lima Giacobbo, B., et al., Brain-Derived Neurotrophic Factor in Brain Disorders: Focus on Neuroinflammation. Mol Neurobiol, 2019. 56(5): p. 3295–3312.

34. Bathina, S. and U.N. Das, Brain-derived neurotrophic factor and its clinical implications. Arch Med Sci, 2015. 11(6): p. 1164–78.

35. Jin, Y., et al., The Role of BDNF in the Neuroimmune Axis Regulation of Mood Disorders. Front Neurol, 2019. 10: p. 515.

36. Paredes, D., A.R. Knippenberg, and D.A. Morilak, Infralimbic BDNF signaling is necessary for the beneficial effects of extinction on set shifting in stressed rats. Neuropsychopharmacology, 2022. 47(2): p. 507–515.

37. Autry, A.E. and L.M. Monteggia, Brain-derived neurotrophic factor and neuropsychiatric disorders. Pharmacol Rev, 2012. 64(2): p. 238–58.

38. Wong, Y.H., et al., Activity-dependent BDNF release via endocytic pathways is regulated by synaptotagmin-6 and complexin. Proc Natl Acad Sci U S A, 2015. 112(32): p. E4475–84.

39. Vermehren-Schmaedick, A., R.A. Khanjian, and A. Balkowiec, Cellular mechanisms of activity-dependent BDNF expression in primary sensory neurons. Neuroscience, 2015. 310: p. 665–73.

40. Miyasaka, Y. and N. Yamamoto, Neuronal Activity Patterns Regulate Brain-Derived Neurotrophic Factor Expression in Cortical Cells via Neuronal Circuits. Front Neurosci, 2021. 15: p. 699583.

41. Kohara, K., et al., Activity-dependent transfer of brain-derived neurotrophic factor to postsynaptic neurons. Science, 2001. 291(5512): p. 2419–23.

42. Horvath, P.M., et al., A subthreshold synaptic mechanism regulating BDNF expression and resting synaptic strength. Cell Rep, 2021. 36(5): p. 109467.

43. Hu, B., A.M. Nikolakopoulou, and S. Cohen-Cory, BDNF stabilizes synapses and maintains the structural complexity of optic axons in vivo. Development, 2005. 132(19): p. 4285–98.

44. Choo, M., et al., Retrograde BDNF to TrkB signaling promotes synapse elimination in the developing cerebellum. Nat Commun, 2017. 8(1): p. 195.

45. Yoshii, A. and M. Constantine-Paton, Postsynaptic BDNF-TrkB signaling in synapse maturation, plasticity, and disease. Dev Neurobiol, 2010. 70(5): p. 304–22.

46. Je, H.S., et al., ProBDNF and mature BDNF as punishment and reward signals for synapse elimination at mouse neuromuscular junctions. J Neurosci, 2013. 33(24): p. 9957–62.

47. Gottmann, K., T. Mittmann, and V. Lessmann, BDNF signaling in the formation, maturation and plasticity of glutamatergic and GABAergic synapses. Exp Brain Res, 2009. 199(3-4): p. 203–34.

48. Bamji, S.X., et al., BDNF mobilizes synaptic vesicles and enhances synapse formation by disrupting cadherin-beta-catenin interactions. J Cell Biol, 2006. 174(2): p. 289–99.

49. Wong-Riley, M.T.T., The critical period: neurochemical and synaptic mechanisms shared by the visual cortex and the brain stem respiratory system. Proc Biol Sci, 2021. 288(1958): p. 20211025.

50. Genoud, C., et al., Altered synapse formation in the adult somatosensory cortex of brain-derived neurotrophic factor heterozygote mice. J Neurosci, 2004. 24(10): p. 2394–400.

51. Shen, K. and C.W. Cowan, Guidance molecules in synapse formation and plasticity. Cold Spring Harb Perspect Biol, 2010. 2(4): p. a001842.

52. Rauti, R., et al., BDNF impact on synaptic dynamics: extra or intracellular long-term release differently regulates cultured hippocampal synapses. Mol Brain, 2020. 13(1): p. 43.

53. West, A.E., Biological functions of activity-dependent transcription revealed. Neuron, 2008. 60(4): p. 523–5.

54. Lu, B., BDNF and activity-dependent synaptic modulation. Learn Mem, 2003. 10(2): p. 86–98.

55. Karpova, N.N., Role of BDNF epigenetics in activity-dependent neuronal plasticity. Neuropharmacology, 2014. 76 Pt C: p. 709–18.

56. Crozier, R.A., et al., BDNF modulation of NMDA receptors is activity dependent. J Neurophysiol, 2008. 100(6): p. 3264–74.

57. Lu, Y., K. Christian, and B. Lu, BDNF: a key regulator for protein synthesis-dependent LTP and long-term memory? Neurobiol Learn Mem, 2008. 89(3): p. 312–23.

58. Panja, D. and C.R. Bramham, BDNF mechanisms in late LTP formation: A synthesis and breakdown. Neuropharmacology, 2014. 76 Pt C: p. 664–76.

59. Aicardi, G., et al., Induction of long-term potentiation and depression is reflected by corresponding changes in secretion of endogenous brain-derived neurotrophic factor. Proc Natl Acad Sci U S A, 2004. 101(44): p. 15788–92.

60. Garad, M., E. Edelmann, and V. Leßmann, Long-term depression at hippocampal mossy fiber-CA3 synapses involves BDNF but is not mediated by p75NTR signaling. Sci Rep, 2021. 11(1): p. 8535.

61. Aarse, J., S. Herlitze, and D. Manahan-Vaughan, The requirement of BDNF for hippocampal synaptic plasticity is experience-dependent. Hippocampus, 2016. 26(6): p. 739–51.

62. Gangarossa, G., et al., BDNF Controls Bidirectional Endocannabinoid Plasticity at Corticostriatal Synapses. Cereb Cortex, 2020. 30(1): p. 197–214.

63. Lu, H., H. Park, and M.M. Poo, Spike-timing-dependent BDNF secretion and synaptic plasticity. Philos Trans R Soc Lond B Biol Sci, 2014. 369(1633): p. 20130132.

64. Leal, G., D. Comprido, and C.B. Duarte, BDNF-induced local protein synthesis and synaptic plasticity. Neuropharmacology, 2014. 76 Pt C: p. 639–56.

65. Gonzalez, M.C., A. Radiske, and M. Cammarota, On the Involvement of BDNF Signaling in Memory Reconsolidation. Front Cell Neurosci, 2019. 13: p. 383.

66. Cunha, C., R. Brambilla, and K.L. Thomas, A simple role for BDNF in learning and memory? Front Mol Neurosci, 2010. 3: p. 1.

67. Bekinschtein, P., M. Cammarota, and J.H. Medina, BDNF and memory processing. Neuropharmacology, 2014. 76 Pt C: p. 677–83.

68. Bekinschtein, P., et al., BDNF is essential to promote persistence of long-term memory storage. Proc Natl Acad Sci U S A, 2008. 105(7): p. 2711–6.

69. Fernandes, D. and A.L. Carvalho, Mechanisms of homeostatic plasticity in the excitatory synapse. J Neurochem, 2016. 139(6): p. 973–996.

70. Rutherford, L.C., S.B. Nelson, and G.G. Turrigiano, BDNF has opposite effects on the quantal amplitude of pyramidal neuron and interneuron excitatory synapses. Neuron, 1998. 21(3): p. 521–30.

71. Leslie, K.R., S.B. Nelson, and G.G. Turrigiano, Postsynaptic depolarization scales quantal amplitude in cortical pyramidal neurons. J Neurosci, 2001. 21(19): p. Rc170.

72. Bolton, M.M., A.J. Pittman, and D.C. Lo, Brain-derived neurotrophic factor differentially regulates excitatory and inhibitory synaptic transmission in hippocampal cultures. J Neurosci, 2000. 20(9): p. 3221–32.

73. Reimers, J.M., J.A. Loweth, and M.E. Wolf, BDNF contributes to both rapid and homeostatic alterations in AMPA receptor surface expression in nucleus accumbens medium spiny neurons. Eur J Neurosci, 2014. 39(7): p. 1159–69.

74. Li, X. and M.E. Wolf, Brain-derived neurotrophic factor rapidly increases AMPA receptor surface expression in rat nucleus accumbens. Eur J Neurosci, 2011. 34(2): p. 190–8.

75. Timmusk, T., et al., Multiple promoters direct tissue-specific expression of the rat BDNF gene. Neuron, 1993. 10(3): p. 475–89.

76. Zheng, F., et al., Regulation of brain-derived neurotrophic factor exon IV transcription through calcium responsive elements in cortical neurons. PLoS One, 2011. 6(12): p. e28441.

77. Sakata, K., et al., Role of activity-dependent BDNF expression in hippocampal-prefrontal cortical regulation of behavioral perseverance. Proc Natl Acad Sci U S A, 2013. 110(37): p. 15103–8.

78. Martinowich, K., et al., Activity-dependent brain-derived neurotrophic factor expression regulates cortistatin-interneurons and sleep behavior. Mol Brain, 2011. 4: p. 11.

79. Ratnu, V.S., W. Wei, and T.W. Bredy, Activation-induced cytidine deaminase regulates activity-dependent BDNF expression in post-mitotic cortical neurons. Eur J Neurosci, 2014. 40(7): p. 3032–9.

80. Hong, E.J., A.E. McCord, and M.E. Greenberg, A biological function for the neuronal activity-dependent component of Bdnf transcription in the development of cortical inhibition. Neuron, 2008. 60(4): p. 610–24.

81. Tuvikene, J., et al., AP-1 Transcription Factors Mediate BDNF-Positive Feedback Loop in Cortical Neurons. J Neurosci, 2016. 36(4): p. 1290–305.

82. Shimada, A., C.A. Mason, and M.E. Morrison, TrkB signaling modulates spine density and morphology independent of dendrite structure in cultured neonatal Purkinje cells. J Neurosci, 1998. 18(21): p. 8559–70.

83. Tongiorgi, E., M. Righi, and A. Cattaneo, Activity-dependent dendritic targeting of BDNF and TrkB mRNAs in hippocampal neurons. J Neurosci, 1997. 17(24): p. 9492–505.

84. An, J.J., et al., Distinct role of long 3’ UTR BDNF mRNA in spine morphology and synaptic plasticity in hippocampal neurons. Cell, 2008. 134(1): p. 175–87.

85. Verpelli, C., et al., Synaptic activity controls dendritic spine morphology by modulating eEF2-dependent BDNF synthesis. J Neurosci, 2010. 30(17): p. 5830–42.

86. Tanaka, J., et al., Protein synthesis and neurotrophin-dependent structural plasticity of single dendritic spines. Science, 2008. 319(5870): p. 1683–7.

87. Pruunsild, P., et al., Dissecting the human BDNF locus: bidirectional transcription, complex splicing, and multiple promoters. Genomics, 2007. 90(3): p. 397–406.

88. Cattaneo, A., et al., The human BDNF gene: peripheral gene expression and protein levels as biomarkers for psychiatric disorders. Transl Psychiatry, 2016. 6(11): p. e958.

89. Crofton, E.J., et al., Topographic transcriptomics of the nucleus accumbens shell: Identification and validation of fatty acid binding protein 5 as target for cocaine addiction. Neuropharmacology, 2021. 183: p. 108398.

90. Zhang, Y., et al., Manipulation of retinoic acid signaling in the nucleus accumbens shell alters rat emotional behavior. Behav Brain Res, 2019. 376: p. 112177.

91. Zhang, Y., et al., Transcriptomics of Environmental Enrichment Reveals a Role for Retinoic Acid Signaling in Addiction. Front Mol Neurosci, 2016. 9: p. 119.

92. Suzuki, K., et al., Convergence of distinct signaling pathways on synaptic scaling to trigger rapid antidepressant action. Cell Rep, 2021. 37(5): p. 109918.

93. Soden, M.E. and L. Chen, Fragile X protein FMRP is required for homeostatic plasticity and regulation of synaptic strength by retinoic acid. J Neurosci, 2010. 30(50): p. 16910–21.

94. Zhang, Y., et al., Rapid single-step induction of functional neurons from human pluripotent stem cells. Neuron, 2013. 78(5): p. 785–98.

95. Zhong, L.R., et al., Retinoic Acid Receptor RARalpha-Dependent Synaptic Signaling Mediates Homeostatic Synaptic Plasticity at the Inhibitory Synapses of Mouse Visual Cortex. J Neurosci, 2018. 38(49): p. 10454–10466.

96. Rios, M., et al., Conditional deletion of brain-derived neurotrophic factor in the postnatal brain leads to obesity and hyperactivity. Mol Endocrinol, 2001. 15(10): p. 1748–57.

97. Luikart, B.W., et al., TrkB has a cell-autonomous role in the establishment of hippocampal Schaffer collateral synapses. J Neurosci, 2005. 25(15): p. 3774–86.

98. Aoto, J., et al., Presynaptic neurexin-3 alternative splicing trans-synaptically controls postsynaptic AMPA receptor trafficking. Cell, 2013. 154(1): p. 75–88.

